# Human genetic variation shapes the antibody repertoire across B cell development

**DOI:** 10.1101/2025.05.19.654982

**Authors:** Oscar L. Rodriguez, Xiang Qiu, Kaitlyn Shields, Christoper Dunn, Amit Singh, Mary Kaileh, Corey T. Watson, Ranjan Sen

## Abstract

The peripheral antibody repertoire is shaped by inherited genetic variation and selection during B cell development. However, how the repertoire changes across developmental stages—and the relative impact of genetics and selection in establishing the antibody repertoire—remain largely unknown. To dissect the individual and combined effects of these factors, we integrated antibody repertoire transcriptomics across pro-B, pre-B, immature, and naive B cells with single-molecule real-time long-read DNA sequencing of the immunoglobulin heavy (IGH), kappa (IGK), and lambda (IGL) chain loci. We find that IGH genetic variants establish gene usage biases at the pro-B stage that persist throughout development. In contrast, IGK and IGL repertoires are extensively remodeled during maturation, consistent with receptor editing. Surprisingly, IGH variants also exert trans effects on light chain gene usage, suggesting coordinated constraints on heavy-light chain pairing. Principal component and allele-specific analyses reveal that genetics primarily shapes the heavy chain repertoire, while selection more strongly influences light chain diversity. These findings define distinct dynamics of antibody repertoire development and underscore the importance of personal immunogenomics in understanding individual immune responses.

## Introduction

The highly diverse human antibody repertoire enables broad protection against a wide range of pathogens ^1–5^. This diversity is generated during B cell development through V(D)J recombination, a somatic process that joins variable (V), diversity (D), and joining (J) genes to form the variable region of the immunoglobulin heavy chain — the portion of the antibody that contributes to antigen binding. V(D)J recombination initiates at the IGH locus in pro-B cells and is followed in pre-B cells by VJ recombination at the IGK or IGL loci to produce the light chains, which pair with the heavy chain to complete the antibody antigen-binding site. Together with junctional diversity and diverse heavy-light chain pairing, this process can generate more than a trillion unique antibody molecules^6,7^. In the bone marrow, the initial antibody repertoire undergoes selection to eliminate autoreactive B cells, ensuring the establishment of a functional and self-tolerant peripheral B cell pool^8^. Early antibody cloning studies demonstrated that approximately 55% to 75% of early B cells exhibit autoreactivity and are either deleted through apoptosis or rescued by receptor editing, which involves replacement of the initial light chain^9^. Supporting this, studies in transgenic mouse models have shown that receptor editing can rescue 25% to 50% of autoreactive B cells through secondary light chain recombination^10,11^. Meanwhile, shallow antibody repertoire sequencing across B cell developmental stages has revealed more pronounced changes in heavy chains compared to light chains, suggesting stronger selection pressures on the heavy chain repertoire^12^.

In addition to selection, inherited genetic variation contributes to antibody repertoire diversity^13–16^. Twin studies have shown that repertoire features are heritable^17,18^, and a recent gene usage QTL study identified genetic variants at the IGH locus—including single nucleotide polymorphisms (SNPs), insertion-deletions (indels), and structural variants (SVs)—associated with inter-individual differences in V, D, and J gene usage in both IgM and IgG repertoires^19^. Critically, the identification of genetic variants influencing gene usage across IGH has been enabled by the application of long-read sequencing. The IGH locus is one of the most structurally complex and polymorphic regions of the human genome^17,18,20–22^, posing major challenges for accurate genotyping^23^. Recent advances in long-read sequencing have enabled near-complete resolution of IGH, IGK, and IGL haplotypes, providing the foundation for high-confidence genetic association analyses across the immunoglobulin loci^24–26^. Together, these findings suggest that both inherited genetic variation and developmental selection shape the antibody repertoire.

However, despite evidence that both selection and genetics shape the antibody repertoire, their respective contributions across B cell development remain poorly defined. Most studies rely on peripheral B cells, limiting the ability to disentangle early genetic effects from changes imposed by selection. In particular, it remains unclear how genetic variation affects repertoire formation at the time of V(D)J recombination, and how this intersects with developmental checkpoints that eliminate autoreactive B cells.

To address this gap, we profiled the IgM, IgD, IgE, IgA, IgG, IgK, and IgL transcriptome across B cell developmental stages—pro-B, pre-B, immature, and naive B cells—in six donors. Using targeted long-read, single-molecule real-time sequencing, we assembled personalized IGH, IGK, and IGL loci, capturing extensive genetic variation, including SNPs and SVs. Using these two paired datasets, we evaluated the antibody repertoire across developmental stages within donors, the impact of genetic variation on the developing repertoire, and the relative contribution of each to establishing the repertoire. We demonstrated that the usage of V, D, and J genes in the heavy and light chain repertoires across B cell subsets is associated with IGH genetic variation, with minimal gene usage changes in the heavy chain repertoire during development and distinct differences in the developmental changes in the kappa and lambda repertoires. These findings reveal that inherited genetic variation predominantly establishes heavy chain repertoire biases at the onset of recombination, while light chain repertoires are more dynamically shaped by developmental selection. Together, our results define distinct roles for genetics and selection in shaping the human antibody repertoire across B cell development.

## Results

### Characterizing the expressed antibody repertoire across B cell development

To profile the antibody repertoire across B cell development, we performed adaptive immune receptor repertoire sequencing (AIRR-seq) of IgM, IgD, IgG, IgE, IgA, IgK, and IgL transcripts using 5’ rapid amplification of cDNA ends (5’ RACE) with UMI barcodes. Libraries were prepared from sorted pro-B, pre-B, and immature B cells from bone marrow, and naive B cells from peripheral blood mononuclear cells (PBMCs) of six healthy donors (D1 – D6) (Figure 1A). B cell subset sorting was validated using AIRR-seq expression profiles. IgM transcripts dominated all B cell stages, while IgD expression increased at the naive B cell stage. Pro-B cells expressed DJ-recombined heavy chain transcripts, consistent with ongoing V(D)J recombination. Light chain transcripts were low at the pro-B stage and primarily corresponded to IGLL1 and IGLL5, encoding the surrogate light chain, with sustained expression in pre-B cells (Supplementary Figures 1–2).

**Figure 1.**
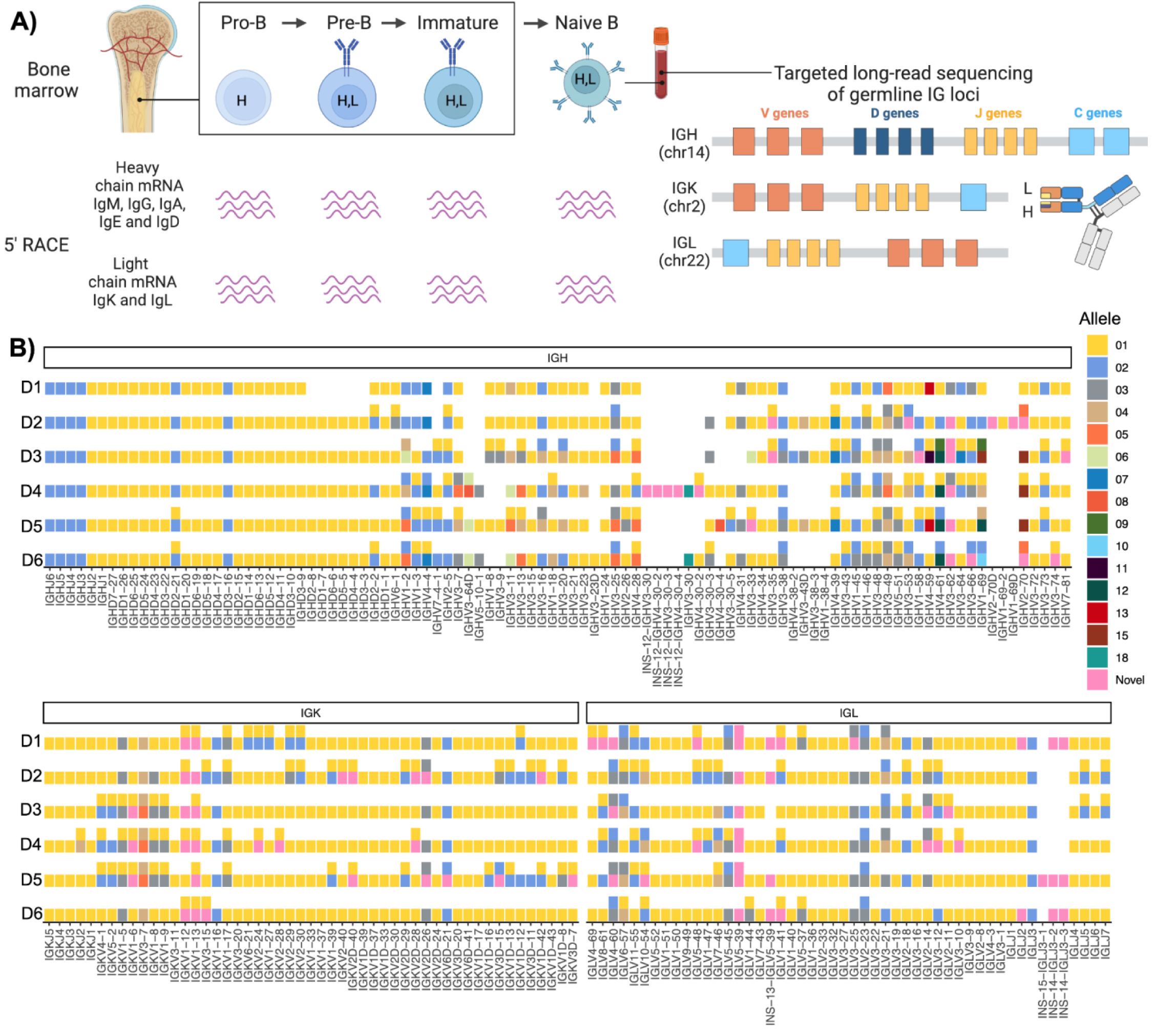
Paired AIRR sequencing of the pro-B, pre-B, immature and naïve heavy, kappa and lambda Ab repertoire and long-read genomic sequencing of the immunoglobulin loci. A) Schematic representation of B cell development, from pro-B to immature B cells in the bone marrow and naive B cells in the periphery. The IGH locus undergoes recombination at the pro-B cell stage (H), while light chain recombination occurs at the pre-B cell stage (L). The antibody repertoire of each B cell subset was analyzed using 5’ RACE to profile the heavy (IgM, IgG, IgA, IgE, and IgD) and light chain (IgK and IgL) repertoires in six individuals. Additionally, the IGK, IGL, and IGH loci were sequenced using targeted long-read sequencing. B) IGH, IGK, and IGL V, D, and J alleles across donors. Blank spaces indicate deleted or missing genes, while pink tiles represent novel alleles not present in IMGT.

A key step in AIRR-seq analysis is the assignment of V, D, and J genes to each sequence. Across the three immunoglobulin loci, there are between 84 and 117 V genes, 23 D genes, and 15–16 J genes, each harboring multiple alleles. The Open Germline Receptor Database (OGRDB) currently curates over 230 IGH V, D, and J alleles^27^. Standard approaches align reads to reference databases such as OGRDB or the ImMunoGeneTics information system (IMGT). However, these methods are limited by missing alleles, the inability to distinguish between deleted and unexpressed genes, and the inability to differentiate duplicated genes encoding identical sequences.

To overcome these challenges, we performed long-read sequencing of each donor’s IGH, IGK, and IGL loci, generating personalized germline allele databases (Figure 1B). To our knowledge, this is the first study to utilize a complete set of personalized germline alleles for AIRR-seq analysis, enabling the most accurate gene and allele assignments to date (Figure 1B). Across donors, we identified an average of 15 novel alleles (absent from IMGT), 16 deleted genes on both maternal and paternal haplotypes, and 25 genes with identical allele sequences within individuals, which were collapsed into composite genes for downstream read assignment.

Each B cell expresses a unique antibody heavy chain, allowing clonal grouping based on identical V, D, and J allele assignments and CDR3 nucleotide sequences. Using IgM AIRR-seq data, we identified an average of 16,760 pro-B, 200,627 pre-B, 119,919 immature, and 254,045 naive heavy chain clones across donors. Light chain clones were grouped based on VJ allele and CDR3 sequence. We identified 9,925 pre-B, 24,793 immature, and 46,992 naive B cell clones in the IGK repertoire, and 2,509 pre-B, 7,691 immature, and 8,090 naive B cell clones in the IGL repertoire. Due to shorter CDR3 regions and the absence of D segments, convergent light chain sequences likely represent multiple B cells.

Together, these data provided the most comprehensive characterization to date of the expressed antibody repertoire across pro-B, pre-B, immature, and naive B cell stages in humans. Compared to previous studies^12^, this dataset represented a ∼24,577% and ∼5,184% increase in the number of analyzed heavy and light chain clones, respectively.

### Light chain gene usage shifts across development, while heavy chain gene usage remains stable

To investigate how the antibody repertoire evolves across B cell development, we quantified V, D, and J gene usage among B cell clones for each donor and B cell subset (Supplementary Figure 3). Overall, IGHV genes exhibited lower variability in usage compared to IGLV (p = 2.4 x 10^-12^) and IGKV (p = 1.1 x 10^-10^) genes across development, as measured by the coefficient of variation and compared using the Wilcoxon rank-sum test (Figure 2A, Supplementary Figure 4A). Among J genes, IGLJ genes showed the highest variability across subsets relative to IGHJ and IGKJ.

**Figure 2.**
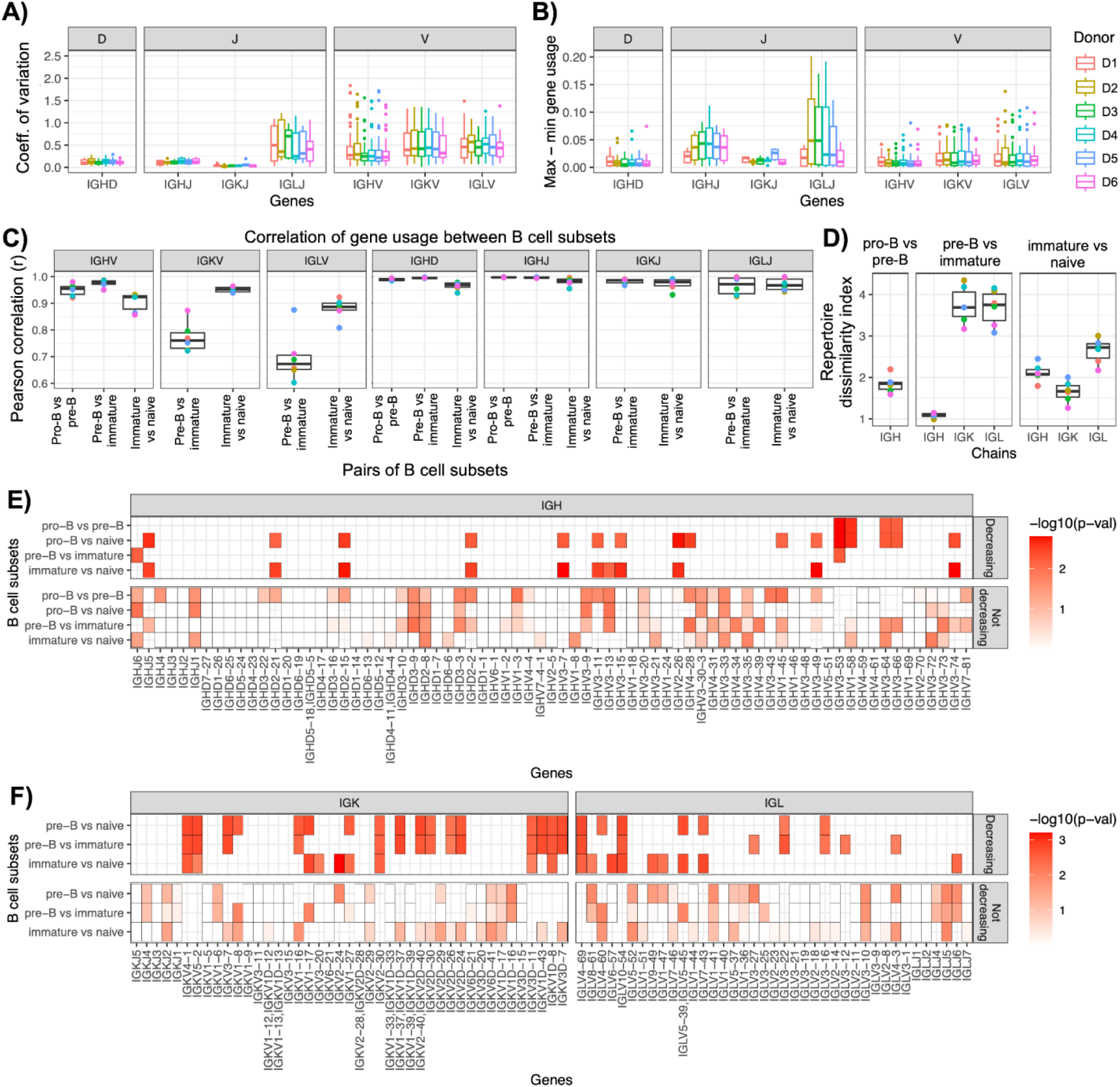
Gene usage dynamics and repertoire remodeling across B cell development. A) Coefficient of variation in V, D, and J gene usage across B cell developmental stages, grouped by donor. B) Difference between maximum and minimum usage frequencies for each gene across B cell development, grouped by donor. C) Pearson correlation of gene usage across donors for consecutive B cell subsets for IGH, IGK, and IGL V, D, and J genes. D) Repertoire dissimilarity index (RDI) measuring differences in VDJ and VJ gene combinations between consecutive B cell subsets within donors. E) IGH genes showing significant decreases in usage across donors between consecutive developmental stages and between the pro-B and naive B cell stages. F) IGK and IGL genes showing significant decreases in usage between consecutive developmental stages and between the pre-B and naive B cell stages.

We next assessed the magnitude of gene usage changes across development by calculating the difference between maximum and minimum usage frequencies for each gene within donors. V gene usage deltas were significantly greater for IGK (p = 1.6 x 10^-8^) and IGL (p = 6.6 x 10^-8^) compared to IGH (Figure 2B, Supplementary Figure 4B), indicating more drastic shifts in light chain repertoires.

To further evaluate gene usage stability across developmental transitions, we calculated Pearson correlation coefficients for gene usage frequencies between consecutive B cell subsets within donors (Figure 2C). IGHV gene usage remained highly stable, with median correlation values of 0.95 (pro-B vs. pre-B), 0.98 (pre-B vs. immature), and 0.92 (immature vs. naive). In contrast, IGKV and IGLV gene usage exhibited greater changes, particularly between the pre-B and immature stages, for which median correlations were 0.67 (IGK) and 0.76 (IGL). J and D gene usage correlations exceeded 0.9 across all comparisons.

We additionally applied the Repertoire Dissimilarity Index^28^ (RDI) to quantify changes in VDJ and VJ gene combinations across developmental stages within donors (Figure 2D). Consistent with correlations observed for V, D, an J segments, individually, RDI values were highest between pre-B and immature stages for IGK and IGL, suggesting that VJ frequencies underwent significant shifts between these stages. RDI analysis also revealed greater changes in IGL gene usage between the immature and naive stages relative to IGK. In contrast, VDJ frequencies for IGH remained relatively stable across all developmental transitions, reflecting minimal gene usage changes in the heavy chain Ab repertoire.

To identify specific genes potentially subject to negative selection during development, we performed paired one-tailed t-tests to detect decreases in gene usage across stages (Figure 2E-F). Across all donors, 16 of 73 (22%) IGH genes showed significant decreases between at least one pair of cell subsets, compared to 14 of 39 (36%) IGL genes and 18 of 43 (42%) IGK genes.

These findings revealed distinct patterns of gene usage dynamics across B cell development. Specifically, whereas IGH gene usage remained relatively stable, IGK and IGL genes underwent more pronounced changes, particularly between the pre-B and immature stages. The differential modulation of V, D, and J gene usage highlighted fundamental differences in the developmental trajectories of the IgM, IgK and IgL antibody repertoires.

### Germline variation establishes early and persistent biases in antibody gene usage across development

Previously, we identified coding and non-coding genetic variants within the IGH locus associated with gene usage (gene usage quantitative trait loci, guQTLs) in the IgM and IgG repertoires of PBMCs. We hypothesized that these genetic effects originate during early B cell development and persist across maturation. To test this, we evaluated guQTL effects across pro-B, pre-B, immature, and naive B cell stages (Figure 3A). Among 37 IGH genes for which previously described guQTLs exhibited genotypic variation in these six donors, 84% (31/37) showed significant associations (p < 0.05) in at least one developmental subset, indicating that genetic influences on gene usage emerge early in B cell development. Notably, for 77% (24/31) of these genes, the genetic associations were consistently detected across all four stages. For example, individuals homozygous for the "G" allele at rs72717212 exhibited higher IGHV1-3 usage compared to carriers of the "T" allele, an effect maintained from the pro-B to naive B cell stages (Figure 3B). Importantly, none of the gene usage-associated IGH SNPs resided within coding regions; 61% mapped to non-coding regions, and the remainder were in linkage disequilibrium with SVs. These findings suggested that germline variation initially modulates gene usage predominantly through regulatory mechanisms that influence V(D)J recombination.

**Figure 3.**
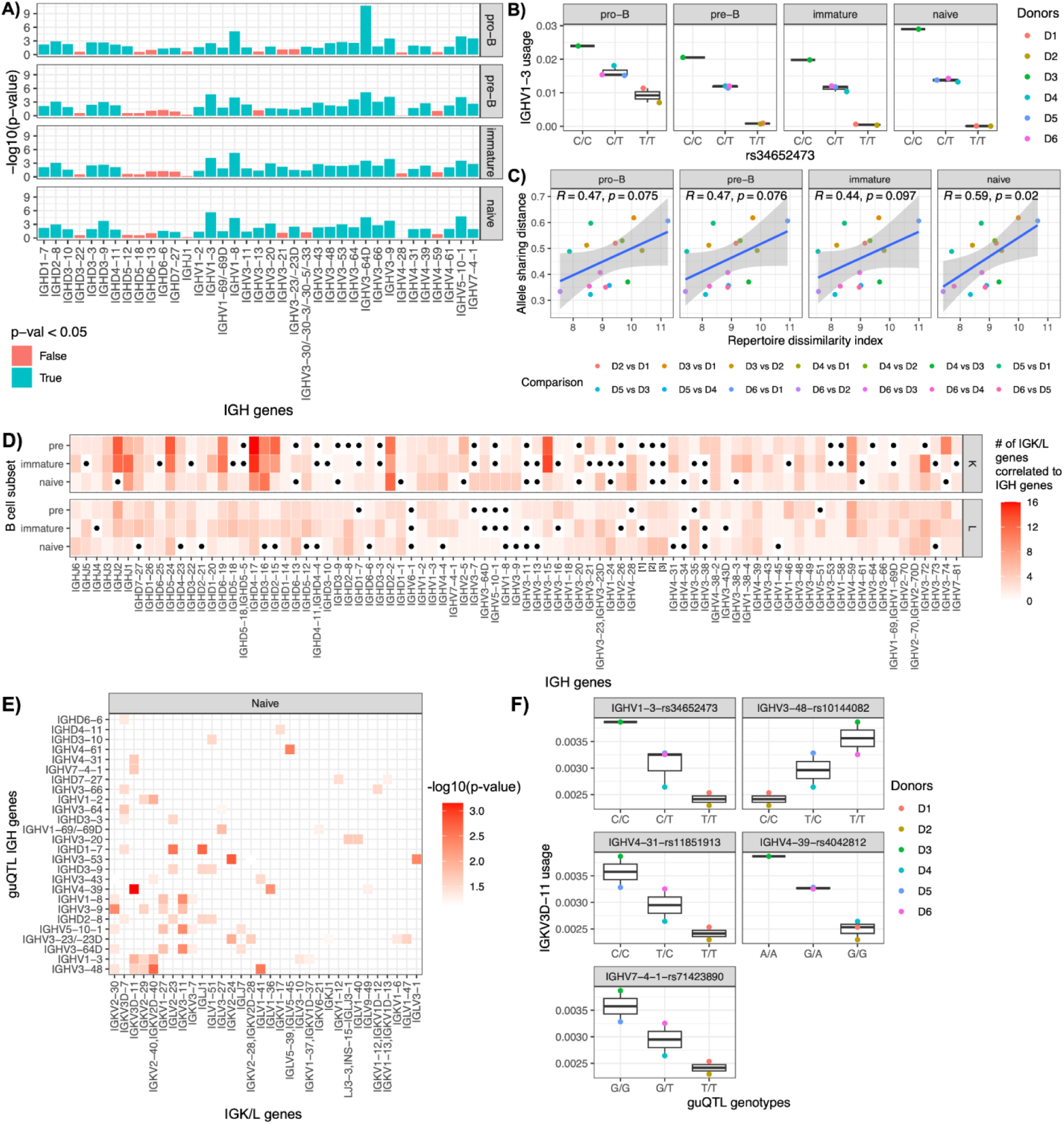
IGH genetic variation impacts heavy and light chain gene usage across B cell development. A) P-values of replicated gene usage QTLs in the pro-B, pre-B, immature, and naive IgM antibody repertoire. B) IGHV1-3 usage in donors D1–D6, stratified by genotypes at the replicated guQTL SNP *rs34652473*. C) Correlation between allele sharing distance (ASD) and repertoire dissimilarity index (RDI) across pro-B, pre-B, immature, and naive IgM repertoires. D) Number of kappa and lambda genes whose usage correlates with that of heavy chain genes. E) P-values for associations between IGH guQTL SNP genotypes and light chain gene usage (IGK and IGL) at the naive B cell stage. F) IGKV3D-11 usage stratified by genotypes at replicated IGH guQTL SNPs associated with *IGHV1-3, IGHV3-48, IGHV4-31, IGHV4-39*, and *IGHV7-4-1* usage.

Given the high rate of guQTL replication across developmental stages, we hypothesized that individuals with more similar IGH haplotypes would have more similar gene usage profiles. To test this, we computed pairwise allele sharing distance (ASD) and RDI between donors. ASD quantified genetic differences at the IGH locus, while RDI measured divergence in VDJ usage frequencies. We found that ASD and RDI were positively correlated, revealing that genetically similar individuals (lower ASD) had more similar repertoire usage profiles (lower RDI) at the pro-B (R = 0.47), pre-B (R = 0.47), and immature (R = 0.44) stages (p < 0.1). This association was strongest at the naive stage (R = 0.59; p = 0.022), highlighting the persistent influence of inherited variation on antibody repertoire composition across B cell development.

### IGH genetic variation exerts *trans* effect on light chain gene usage

During B cell development, heavy chain V(D)J recombination precedes light chain V(D)J recombination, in that light chains are first expressed at the pre-B cell stage and paired with pre-formed heavy chains. This sequential process raises the potential of dependencies between heavy and light chain gene usage. To test this, we assessed correlations between heavy and light chain gene usage across donors (Figure 3D). Strikingly, all 85 heavy chain genes exhibited significant correlations (FDR < 0.05) with at least one light chain gene in at least one B cell subset. In naive B cells, each heavy chain gene was correlated, on average, with two IGK genes and one IGL gene. Notably, IGHD4-17 had the highest number of statistically significant correlations with IGK genes (n = 16) at the pre-B stage. IGHV4-59 had the most correlations with IGL genes (n = 7) at the same stage.

Given the strong genetic influence on heavy chain V(D)J recombination (Figure 3A–C) and the observed heavy-light chain correlations (Figure 3D), we hypothesized that IGH genetic variants might indirectly modulate light chain gene usage. To test this, we assessed whether lead IGH guQTL SNPs were associated with usage of light chain genes correlated with their respective IGH genes. This analysis revealed that 28 of 31 lead guQTL SNPs were significantly associated with the usage of 59 IGK or IGL genes in at least one B cell subset (Figure 3E). For example, lead guQTLs for IGHV1-3, IGHV3-48, IGHV4-31, IGHV4-39, and IGHV7-4-1 were also associated with IGKV3D-11 usage in the naive repertoire (Figure 3F).

Together, these findings suggested that germline variation within the IGH locus not only shapes heavy chain gene usage but also exerts *trans* effects on light chain gene usage during B cell development. In this context, *trans* effects refer to instances where genetic variation at one genomic locus (e.g., IGH) influences the recombination or expression frequency of genes located at distinct, unlinked loci (e.g., IGK or IGL). These results indicated that inherited variation in the heavy chain region may indirectly influence light chain gene usage, contributing to inter-individual diversity in antibody repertoires. This highlights an additional layer of germline impact on antibody repertoire formation, extending beyond *cis* effects at the IGH locus to *trans* effects across immunoglobulin loci.

### Genetic background predominantly shapes heavy chain repertoires, while selection refines light chains

Results presented above have demonstrated that both germline variation (Figure 3A–C) and B cell development (Figure 2E–F) can exert influence over the composition of the repertoire. We next investigated the relative contributions of genetic background and developmental selection in shaping antibody repertoires. We reasoned that if selection dominated repertoire formation, gene usage profiles within repertoires of the same B cell subsets across individuals would be more similar, whereas if genetic effects were to dominate, we would expect that repertoires from different B subsets within a donor would be more similar to one another than to their counterparts in other donors. To assess this, we performed principal component analysis (PCA) of gene usage across all donors and B cell subsets (Figure 4A). Consistent with strong genetic control, PCA revealed that for IGH, within-donor distances between B cell subsets were significantly smaller than between donor distances of the same B cell subsets (p = 0.0009) (Figure 4B). In contrast, for IGK and IGL, within-donor distances were significantly larger, indicating a greater role for selection in shaping light chain repertoires.

**Figure 4.**
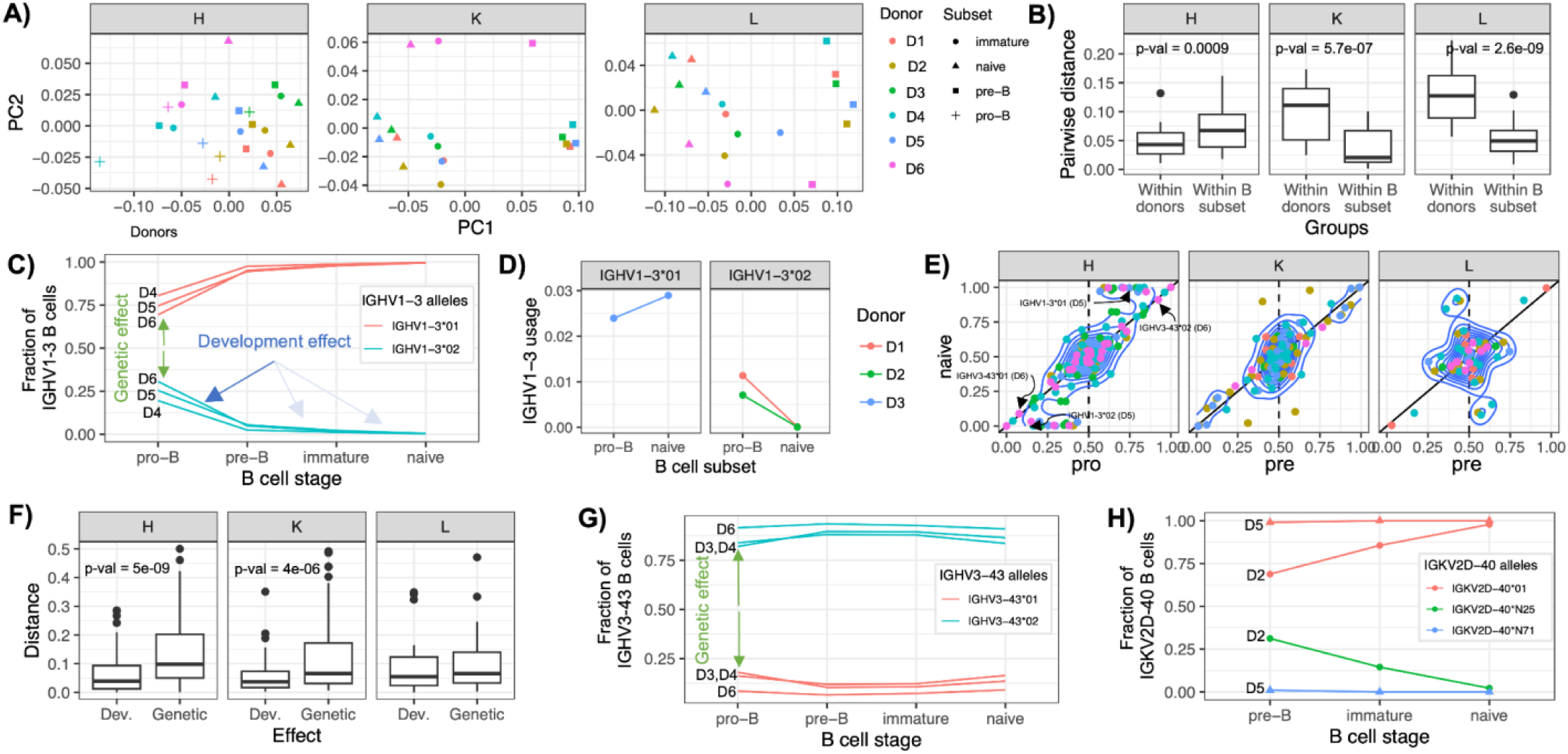
Genetic and development effects on the developing Ab repertoire. A) Principal component analysis (PCA, unscaled) of IGH, IGK, and IGL V gene usage across all donors and B cell subsets. B) Distribution of PCA-based distances from (A) between B cell subsets within donors and between subsets across donors for IgM, IgK, and IgL repertoires. C) Frequency of IGHV1-3*01 and IGHV1-3*02 alleles among IGHV1-3-expressing B cells across B cell development. D) IGHV1-3 usage in the pro-B and naive antibody repertoires of donors homozygous for IGHV1-3*01 or IGHV1-3*02. E) Allele frequencies of heterozygous V genes in the pro-B (or pre-B) and naive B cell stages. Solid line represents y = x (no selection); dotted line represents a pro/pre-B allele frequency of 0.5 (no genetic bias). F) Distribution of allele deviations relative to the y = x line (selection effects) and to the vertical line at x = 0.5 (genetic effects), based on (E). G) Frequency of IGHV3-43*01 and IGHV3-43*02 alleles among IGHV3-43-expressing B cells across B cell development. H) Frequency of IGK2D-40*01, IGK2D-40*N25, and IGK2D-40*N71 alleles among IGK2D-40-expressing B cells across B cell development.

We next sought to investigate the extent of genetic and developmental effects at the gene and allele level. To do this, we identified heterozygous IG genes across individuals, for which we could separately quantify differential genetic and selective effects of each allele (Figure 4C). For example, donors D4–D6, were heterozygous for IGHV1-3, carrying both IGHV1-3*01 and IGHV1-3*02 alleles. In the pro-B cells of these individuals, we found that IGHV1-3 BCRs were predominantly represented by the *01 allele (allele fraction among IGHV1-3 BCRs: D4, 0.8; D5, 0.75; D6, 0.73). This bias indicated differential allele usage, which could be directly attributed to a genetic effect. However, this bias was amplified by selection, in that over the course of development, the relative proportion of IGHV1-3*02 further decreased from pre-B to naive B cells (Figure 4D). This trend was directly observable in the context of IGHV1-3 guQTL genotypes (Figure 3B), in that donors D1 and D2, who were homozygous for IGHV1-3*02, exhibited the lowest IGHV1-3 usage in the pro-B repertoire compared to donors of other guQTL genotypes (Figure 3B). Notably, although the usage biases observed in pro-B cells were attributable to genetics, additional effects of selection became apparent in the other B cell subsets, as we observed that IGHV1-3*02 usage in D1 and D2 declined to zero by the naive B cell stage (Figure 4D).

To systematically quantify the relative gene-level contributions of genetics and selection across the repertoire, we compared allele frequency proportions (as shown for IGHV1-3) at all heterozygous genes in pro-B (for heavy chains) and pre-B (for light chains) to those in naive B cells (Figure 4E). We defined the genetic effect as the deviation of initial allele frequencies from 0.5 at the pro-B (for IGH genes) and pre-B cell (for IGK/L genes) stages; this assumed that 0.5 represented the expected gene-level allele frequency if no allele-specific biases in V(D)J recombination were present. A substantial deviation from 0.5 at the pro-B or pre-B stage indicates a genetic bias affecting allele-specific usage. The selection effect was quantified by comparing allele frequencies between early (pro-B or pre-B) and late (naive) stages. In the absence of selection, allele frequencies would remain stable across B cell subsets, falling along the y = x line when early and late frequencies are plotted against one another. In contrast, deviations from this line would reflect the action of selection, favoring or disfavoring particular alleles during B cell maturation. Using this approach, we found that among heterozygous IGHV genes, genetic effects were more dominant, evidenced by substantial initial deviations from 0.5, compared to relatively smaller shifts observed in allele usage between cell subsets (Figure 4F). Surprisingly, a similar pattern was observed for IGKV genes. However, for IGLV genes, we noticed a contrasting pattern, in which allele usage changes between cell types were larger than those initially observed in the pre-B subset; this was consistent with observations made by PCA (Figure 4B).

Finally, this allele-specific analysis revealed distinct patterns, including cases where genetic effects were observed without detectable selection, and instances of complete allelic silencing (Figure 4H). For example, the novel allele IGKV2D-40*N25 was completely absent among IGKV2D-40–expressing pre-B cells in donor D5, suggesting a lack of functional rearrangement or expression (Figure 4I). Thus, while we observe evidence that selection is more broadly influencing K and L expression across the repertoire as a whole, genetics may still exert stronger effects than development for specific genes.

## Discussion

By integrating matched antibody repertoire transcriptomics across B cell developmental stages with long-read sequencing of the IGH, IGK, and IGL loci, we delineate the distinct and combined roles of germline predisposition and developmental selection in establishing peripheral antibody diversity.

Our results demonstrate that genetic variation within the IGH locus significantly influences antibody gene usage beginning at the pro-B cell stage, when V(D)J recombination is initiated. These genetic effects persist throughout B cell development resulting in gene usage profiles that are maintained across developmental transitions. This persistence suggests that IGH genetic variation is not simply a modifier of the peripheral antibody repertoire but a foundational determinant of its composition, with recombination biases established early and carried forward through selection.

Importantly, the guQTL variants were exclusively in non-coding regions. Our prior work showed that these variants are enriched in CTCF binding sites and distal enhancer sites. In mouse models, chromatin state and three-dimensional genome organization are known to control V(D)J recombination efficiency and accessibility^29–32^. Together, these findings support a model in which non-coding IGH variants influence the regulation of V(D)J recombination, likely through modulation of the local epigenetic or chromatin landscape. Such regulatory differences may underlie inter-individual variation in baseline antibody repertoire composition and, ultimately, immune response potential.

While the absence of guQTLs in the light chain loci limits our ability to assess the impact of IGK and IGL genetic variation on light chain gene usage, our data reveal *trans*-effects in which IGH genetic variation influences light chain repertoire composition. These findings suggest that IGH variants may shape light chain selection indirectly, potentially through pairing constraints that favor specific heavy-light chain combinations or modulate B cell fitness. This interlocus influence highlights the coordinated nature of antibody repertoire formation and underscores the need to consider both chains together. Future work incorporating paired-chain or single-cell repertoire data will be essential to disentangle *trans*-acting effects from local cis-regulatory mechanisms at the light chain loci.

In contrast to the relatively stable heavy chain repertoire, we observed substantial shifts in light chain gene usage across development—particularly between the pre-B and immature B cell stages—consistent with strong selection acting on the light chain. These changes are likely driven by receptor editing, a mechanism that allows B cells to replace autoreactive light chains while preserving productive heavy chains. The absence of corresponding shifts in the IGH repertoire supports this interpretation, as widespread clonal deletion would be expected to impact both chains. Our findings are consistent with knock-in mouse models in which receptor editing was the primary mechanism for rescuing autoreactive B cells while minimizing loss of functional heavy chains^33^. Together, these results support a developmental division of labor: germline variation establishes heavy chain usage biases early in B cell development, while light chain repertoires are refined by selection to enforce self - tolerance and optimize antigen recognition^8^.

Understanding how inherited genetic variation shapes the developing antibody repertoire has important implications for both basic immunology and personalized medicine^34^. Differences in IGH, IGK, and IGL haplotypes likely contribute to individual variability in immune responses to pathogens, vaccines, and autoimmune triggers. Because V, D, and J gene usage influences the properties of antibody binding regions, early biases in recombination may affect the probability of generating effective or autoreactive antibodies. Our findings suggest that integrating personal germline genotypes with repertoire data could enable predictive models of immune competence, vaccine responsiveness, and disease susceptibility^19^. These insights may also inform the rational design of monoclonal antibodies and vaccines^35^ optimized for individuals with specific immunogenetic backgrounds.

In summary, our study provides a comprehensive view of how genetic and developmental forces shape the human antibody repertoire. Germline variation in the IGH locus establishes early and persistent gene usage biases, while receptor editing sculpts the light chain repertoire during key developmental transitions. The identification of *trans*-effects between IGH and light chain loci adds a new dimension to our understanding of antibody generation and highlights the need to consider both genetic architecture and developmental context in studies of immune diversity. These findings lay the groundwork for future research aimed at uncovering the molecular mechanisms linking genotype to function and advancing the clinical application of immunogenomic insights.

### Limitations of study

Our study has limitations that should be acknowledged. The sample size of six individuals, while providing deep insights, does not capture the full extent of genetic diversity present in broader populations. Larger cohorts are necessary to validate our findings and explore the influence of rare genetic variants. Additionally, our analysis focused on the unpaired heavy and light chain repertoires; pairing information could provide more detailed insights into BCR specificity and functionality. Future studies should aim to include more diverse populations to assess the generalizability of our findings. Integrating single-cell sequencing technologies could also allow for the analysis of paired heavy and light chains, providing a more complete picture of BCR diversity and its functional implications. Moreover, investigating how environmental factors, such as exposure to different pathogens, interact with genetic determinants to shape the Ab repertoire would be valuable.

## Methods

### Isolation of mononuclear cells from human blood and bone marrow

Donor matched blood (50 ml) and bone marrow (25 ml) were isolated with Ficoll-Paque PLUS (Cytiva, Cat# 17144002) according to Miltenyi Biotec protocol. Carefµlly layered 35 ml of diluted blood or bone marrow over 15 ml of Ficoll-Paque PLUS in a 50 ml conical tube. Centrifuged at 445xg for 35 minutes at 20 °C in a swinging bucket rotor without brake. Aspirated the upper layer leaving the mononuclear cell layer undisturbed at the interphase. Carefµlly transferred the blood or bone marrow mononuclear cells at the interphase to a new 50 ml conical tube. Washed cells by adding up to 40 ml of buffer, mixed gently and centrifuged at 300xg for 10 minutes at 20 °C. Carefµlly remove supernatant completely. For removal of platelets, resuspended the cell pellet in 50 ml of stain buffer (FBS) (BD, Cat# 554656) and centrifuged at 200xg for 10 minutes at 20 ^0^C. Carefµlly removed the supernatant completely. Resuspended cell pellet in ice cold stain buffer (FBS) (BD, Cat# 554656) with concentration of 1X10^8^/100 µl buffer.

### Antibodies label and FACS sort

Mononuclear cells from blood were labelled with anti-human CD19 (2.5 µl /100 µl), anti-human CD27 (2.5 µl /100 µl), anti-human CD38 (2.5 µl /100 µl) and DRAQ7 (4 µl of 100 fold diluted /100 µl) on ice for 30 minutes. Mononuclear cells from bone marrow were labelled with anti-human CD19 (2.5 µl /100 µl), anti-human CD34 (2.5 µl /100 µl), anti-human CD45 (2.5 µl /100 µl), anti-human IgM (1.25 µl /100 µl), anti-human IgD (1.25 µl /100 µl), anti-human CD10 (2.5 µl /100 µl) and DRAQ7 (4 µl of 100 fold diluted/100 µl) on ice for 30 minutes. Cells were washed with ice cold stain buffer (FBS) twice. Blood derived human naïve B cells were sorted as DRAQ7^-^ CD19^+^CD38^+^CD27^-^. Bone marrow derived human pro-B, pre-B and immature B cells were sorted as DRAQ7^-^ CD45^+^CD10^+^CD34^+^CD19^+^IgM^-^IgD^-^, DRAQ7^-^CD45^+^CD10^+^CD34^-^CD19^+^IgM^-^IgD^-^ and DRAQ7^-^CD45^+^CD10^+^CD34^-^ CD19^+^IgM^+^IgD^-^, respectively.

### RNA extraction

Cells were suspended in 300-600 µl TRIzol reagent (Thermofisher, Cat# 15596026). Genomic DNA free RNA was extracted with Direct-zol RNA Microprep Kits (Zymo Research, Cat# R2062) according to manufacturer protocol. Added an equal volume ethanol (95-100%) (300-600 µl) to a sample lysed in TRIzol reagent and mixed thoroughly. Transferred the mixture into a Zymo-Spin™ IC Column in a collection tube and centrifuged. Transferred the column into a new collection tube and discarded the flow-through. Added 400 µl RNA Wash Buffer to the column and centrifuge. In an RNase-free tube, added 5 µl DNase I (6 U/µl), 35 µl DNA Digestion Buffer and mixed by gentle inversion. Added the mix directly to the column matrix. Incubated at room temperature (20-30°C) for 15 minutes. Added 400 µl Direct-zol™ RNA PreWash to the column and centrifuge. Discarded the flow-through and repeated this step. Added 700 µl RNA Wash Buffer to the column and centrifuge for 1 min to ensure complete removal of the wash buffer. Transferred the column carefully into an RNase-free tube. To elute RNA, added 15 µl of DNase/RNase-Free Water directly to the column matrix and centrifuged. RNA was quantified with Qubit RNA HS (High Sensitivity) Assay Kit (Thermofisher, Cat# Q32855).

### Human BCR repertoire generation

Human BCR repertoire were generated with SMART-Seq® Human BCR (with UMIs) (TakaRa, Cat# 634778) according to manufacturer protocol. Thawed the First-Strand Buffer at room temperature. Did not store on ice. Thawed BCR Enhancer, SMART UMI Oligo, and dT Primer on ice. Gently vortexed each reagent to mix and centrifuged briefly. Stored on ice until needed. Preheated the thermal cycler to 72°C. On ice, prepared samples and controls in nuclease-free, thin-wall PCR tubes, plates, or strips by adding the reagents in the order shown below: Sample (100ng), Nuclease-Free Water (up to 8.5 µl), BCR Enhancer (1 µl), dT Primer (2 µl). Mixed by gently vortexing and then centrifuged briefly. Incubated the tubes or plates at 72°C in the preheated, heated-lid thermal cycler for 3 min. During this incubation, prepared the RT Master Mix. At room temperature, prepared RT Master Mix by combining the following components in the order shown in the table below. Mixed the RT Master Mix well by gently pipetting up and down then centrifuge briefly. RT Master Mix: First-Strand Buffer (1 µl), SMART UMI Oligo (1 μl), RNase Inhibitor (0.5 μl), SMARTScribe Reverse Transcriptase (2 µl). Immediately after the 3 min incubation at 72°C (Step 6), placed the samples on ice for 2 min. Removed the SMARTScribe Reverse Transcriptase from the freezer, centrifuged briefly, and stored on ice. Reduced the temperature of the thermal cycler to 42°C. Added the SMARTScribe Reverse Transcriptase to the RT Master Mix according to the table above. Mixed well by gently pipetting up and down. Added 7.5 μl of the RT Master Mix to each reaction tube or well. Mixed the contents of each tube or well by pipetting gently and centrifuged briefly. Placed the tubes/plate in the preheated thermal cycler and run the following program: 42°C (90 min), 70°C (10 min), 4°C (forever). Thawed all the reagents needed for PCR on ice. Gently vortexed each reagent tube (except PrimeSTAR GXL Premix) briefly to mix and spined down. Stored on ice. Prepared PCR1 Master Mix by combining the following components in the order shown in the table below. Gently vortexed to mix then briefly centrifuged. PCR1 Master Mix: Nuclease-Free Water (3 μl), hBCR PCR1 Universal Forward (1 μl), hBCR PCR1 Reverse (1 μl) and PrimeSTAR GXL Premix (25 μl). Added 30 μl of the PCR1 Master Mix to each tube or well containing 20 µl of the first-strand cDNA produced from above. Mixed well and briefly spined to collect the contents at the bottom of the tubes/wells. Placed the tubes/plate in a preheated thermal cycler with a heated lid and run the following program: 95°C (1 min), 16 cycles-98°C (10 sec)- 60°C (15 sec)- 68°C (45 sec), 4°C (forever). Thawed all the reagents needed for PCR on ice. Gently vortexed each reagent tube (except PrimeSTAR GXL Premix) briefly to mix and spined down. Stored on ice. On ice, prepared a PCR 2 Master Mix by combining the following components in the order shown in the table. Gently vortexed to mix and briefly centrifuged. PCR 2 Master Mix: Nuclease-Free Water (21 μl), hBCR PCR2 HC Reverse -or hBCR PCR2 LC Reverse (1 μl), PrimeSTAR GXL Premix (25 μl). For each reaction, added 47 μl of PCR 2 Master Mix to nuclease-free, thin-wall, 0.2 ml PCR tubes or plate wells. Added 1 μl of appropriate PCR1 product to each corresponding PCR 2 tube/well. Add 2 μl of the appropriate UDI (12.5 µM) to each tube/well. Gently vortexed to mix and briefly spined to collect the contents at the bottom of the tubes/wells. Placed the tubes/plate in a preheated thermal cycler and run the following program: 95°C (1 min), 18 cycles-98°C (10 sec)-60°C (15 sec)-68°C (45 sec), 4°C (forever).

### Human BCR repertoire purification and sequencing

Human BCR repertoire purification was performed according to Takara protocol (Cat# 634778). Vortexed SPRI beads until evenly mixed, then added 30 μl of the beads to each sample (50 μl). Mixed thoroughly by gently pipetting the entire volume up and down at least 10 times. Incubated at room temperature for 8 min to let the DNA bind to the beads. Briefly spined the samples to collect the liquid from the side of the tube or sample well. Placed the samples on the magnetic separation device for ∼5 min or longer until the liquid appeared completely clear, and there were no beads left in the supernatant. While the reaction tubes were sitting on the magnetic separation device, used a pipette to transfer the supernatant (which contains your library) to clean PCR tubes. After transferring, removed the tubes containing the beads from the magnetic separation device and discarded them. Added 30 μl of beads to each tube containing supernatant. Mixed thoroughly by gently pipetting the entire volume up and down at least 10 times. Incubated at room temperature for 8 min to let the DNA bind to the beads. Placed the tubes on the magnetic separation device for ∼10 min or until the solution was completely clear. The libraries were now bound to the beads. While the tubes were sitting on the magnetic separation device, removed the supernatant with a pipette and discarded it. Kept the tubes on the magnetic separation device. Added 200 μl of freshly made 80% ethanol to each sample without disturbing the beads, to wash away contaminants. Waited for 30 sec and used a pipette to carefully remove the supernatant containing contaminants. The library remained bound to the beads during the washing process. Repeated the ethanol wash once more. Briefly spined the tubes (∼2,000g) to collect the remaining liquid at the bottom of each tube. Placed the tubes on the magnetic separation device for 30 sec, then removed all remaining liquid with a pipette. Let the sample tubes rest open on the magnetic separation device at room temperature for ∼2–2.5 min until the pellet appeared dry and was no longer shiny. Once the bead pellet has dried, removed the tubes from the magnetic separation device and added 15 μl of Elution Buffer to cover the pellet. Mixed thoroughly by pipetting up and down to ensure complete bead dispersion. Incubated at room temperature for at least 5 min to rehydrate. Briefly spined the samples to collect the liquid from the side of the tube or sample well. Placed the samples back on the magnetic separation device for 2 min or longer until the solution was completely clear. Transferred clear supernatant containing purified BCR library from each tube to a nuclease-free, low adhesion tube. Labeled each tube with sample information and store at –20°C.

Pooled heavy chain libraries were sequenced using NextSeq 2000 with 2 × 300 bp. Each individual library was sequenced with an average depth of 5 million reads. Pooled heavy chain libraries were also sequenced using NovaSeq 6000 SP with 2 × 300 bp. Each individual library was sequenced with an average depth of 15 million reads.

### Long-read library preparation, sequencing and processing

IGH, IGK, and IGL loci were sequenced using the Pacific Biosciences Sequel IIe platform following previously described methods^19,24–26^. Briefly, high molecular weight genomic DNA was extracted from PBMCs. For targeted enrichment of the IGH, IGK, and IGL loci, a combined Roche HyperCap DNA probe panel was utilized. Genomic DNA (0.5–2 μg) was sheared to ∼8–16 kbp using g-tubes (Covaris, Woburn, MA, USA) and size-selected using the Blue Pippin system (Sage Science, Beverly, MA, USA), collecting fragments between 5–9 kbp. DNA was end-repaired and A-tailed using the KAPA HyperPrep protocol (Roche), ligated to universal adapters, and PCR-amplified. Samples were pooled and subjected to a single combined capture using the custom HyperCap probe panel. After hybridization and capture, libraries were PCR-amplified to enrich targeted fragments. Enriched libraries were prepared for SMRT sequencing using the SMRTbell Express Template Preparation Kit 2.0 according to the manufacturer’s protocol. Sequencing was performed as multiplexed pools (12-plex to 24-plex) using Sequel IIe 2.0 chemistry with 30-hour movies. The generated long-read sequencing data were processed using IGenotyper^22^ for haplotype-resolved IG locus variant and allele identification. Structural variants (SVs) were detected using assemblies generated with hifiasm^36^.

### Processing AIRR-sequencing data

Paired-end AIRR-seq reads were processed using the pRESTO toolkit^37^. Reads were filtered to retain only those with a minimum Phred quality score of 20 using FilterSeq.py quality -q 20. Primer sequences were removed with MaskPrimers.py using the parameters --start 7 --mode cut for Read 1 and --start 12 --mode cut --barcode for Read 2, preserving the unique molecular identifier (UMI) barcode from Read 2. The barcode was appended to Read 1 using PairSeq.py --2f BARCODE, and reads with matching barcodes were aligned using AlignSets.py muscle --bf BARCODE. Barcode consensus sequences were generated with BuildConsensus.py --prcons 0.6 --maxerror 0.1 -n 1. Following consensus building, paired reads were sorted and assembled into full-length sequences using PairSeq.py and AssemblePairs.py align. Assembled sequences were collapsed to remove duplicates using CollapseSeq.py, and only sequences supported by more than two contributing reads were retained using SplitSeq.py group --num 2. Personalized IGH, IGK, and IGL germline allele databases were constructed from long-read assemblies using receptor_utils and NCBI’s makeblastdb tool. The make_igblast_ndm and annotate_j utilities were used to generate IMGT-gapped V-region sequences and auxiliary J files. Personalized BLAST databases were built separately for IGH, IGK, and IGL loci. Processed IgM, IgK, and IgL AIRR-seq sequences were aligned to their respective personalized germline databases using igblastn with the following parameters: -outfmt 19, -ig_seqtype Ig, -domain_system imgt, and -organism human.

### Clonal assignment using AIRR-sequencing data

The Hamming distance between IgM sequences was calculated using the distToNearest function from the shazam package. Clones were assigned with DefineClones.py using the parameters --act set, --model ham, -- norm len, and --dist 0.05. The resulting Change-O file was filtered to retain only productive sequences with assigned V and J genes (v_call and j_call), 100% V-gene identity (v_identity = 100%), and a single representative clone per B cell stage.Light chain repertoires (IgK and IgL) were processed similarly; however, instead of defining clonal groups, a unique B cell was defined as a distinct combination of v_call, j_call, and junction sequence. One representative sequence per B cell was retained for downstream analyses. Gene usage analyses were performed by counting the number of clones (for IgM) or unique B cells (for IgK and IgL) assigned to each V, D, and J gene. Frequencies were calculated by normalizing counts to the total number of clones or cells within each donor and B cell stage.

### Replicating of gene usage QTLs

GuQTLs were obtained from Rodriguez et al. (2023)^19^. For each previously identified SNP-gene pair, we assessed whether the six donors included at least one homozygous reference, one heterozygous, and one homozygous alternate genotype. SNPs meeting this criterion were tested for association with gene usage in our dataset using a linear regression model. A guQTL was considered replicated if the association p-value was less than 0.05.

## Resource availability

Requests for further information and resources should be directed to and will be fulfilled by the lead contact, Ranjan Sen.

## Acknowledgments

We thank all the participants who kindly donate bone marrow and blood samples for research purposes. This research was supported by the Intramural Research Program of the National Institute on Aging.

## Author contributions

O.L.R., X.Q., C.T.W., and R.S. contributed to experimental design and discussions. X.Q., C.D., A.S., and M.L. performed flow sorting and AIRR sequencing experiments. K.S. prepared samples for long-read sequencing. O.L.R. performed AIRR-seq and genetic analyses. O.L.R., X.Q., C.T.W., and R.S. interpreted the data and results, and wrote the manuscript.

